# Single-cell transcriptomic profiling reveals a novel signature of necrotizing granulomatous lesions in the lungs of *Mycobacterium tuberculosis*-infected C3HeB/FeJ mice

**DOI:** 10.1101/2025.05.06.652098

**Authors:** Shintaro Seto, Shiho Ohmori, Hajime Nakamura, Minako Hijikata, Naoto Keicho

## Abstract

Tuberculosis (TB) pathology involves complex immune responses within granulomatous lesions. Using single-cell RNA sequencing, we characterized the cellular compositions of necrotizing granulomatous lesions that developed in the lungs of *Mycobacterium tuberculosis*-infected C3HeB/FeJ mice. We identified 11 distinct major cell types, including phagocytes such as neutrophils and macrophages, and T cells, natural killer cells, B cells, dendritic cells, and plasmacytoid dendritic cells. Among T cells, particularly, *Pdcd1⁺* γδ T cells were detected in necrotizing granulomatous lesions, suggesting their potential role in the pathogenicity of *M. tuberculosis*. Within the macrophage populations, we identified a cluster with significantly higher *Plin2* expression compared to other clusters, whose transcriptomic profile was consistent with that of foamy macrophages. A subset of the *Plin2*-expressing macrophages was identified as a major source of *Ifnb1* and *Cxcl1*, suggesting their involvement in type I interferon signaling and neutrophil recruitment. Furthermore, we identified *Flrt2*, *Hyal1*, and *Mmp13* as novel molecular markers of *Plin2*-expressing macrophages, which were localized to the peripheral rim regions of necrotizing granulomas. In conclusion, our results provide the immune landscape of necrotizing granulomas and reveal novel functional states of macrophages contributing to TB pathogenesis.

## Introduction

*Mycobacterium tuberculosis* infects approximately a quarter of the global population and remains the causative agent of tuberculosis (TB), which is one of the leading causes of death worldwide (1). Over a lifetime, 5–10% of infected individuals eventually develop active TB disease. Understanding the immunological conditions leading to TB progression is crucial for the development of new vaccines, early diagnostics, and host-directed therapies.

Upon inhalation, *M. tuberculosis* bacilli are phagocytosed by alveolar macrophages, followed by their migration into the interstitial space in the lungs (2). Due to their inability to control the intracellular replication of *M. tuberculosis* (3), infected macrophages secrete cytokines and chemokines that recruit lymphocytes and additional macrophages from blood vessels. The resulting aggregation of immune cells leads to the formation of granulomas (4). Despite the heterogeneity of granulomatous lesions, necrotizing granulomas are a pathological hallmark in TB patients (5–7). Moreover, foamy macrophages play critical roles in granuloma formation, development, maintenance, and dissemination of infection (5, 8). These foamy macrophages serve as a niche for *M. tuberculosis* replication (9), and their cell death is believed to contribute to the formation of necrotic cores within granulomas (5, 8).

Recent studies on TB pathology have focused on the cellular heterogeneity of granulomas by analyzing their cellular composition and transcriptomic profiles. Advanced technologies such as single-cell RNA sequencing (scRNA-seq) and spatial transcriptomics have been applied to lung tissues from TB patients, and *M. tuberculosis*-infected non-human primates and mice (10–28). Although transcriptomic analyses combined with laser microdissection have revealed significantly elevated *Plin2* expression in foamy macrophages within necrotizing granulomas in the lungs of both TB patients and C3HeB/FeJ mice (29, 30), single-cell transcriptomic profiling specifically targeting foamy macrophages remains limited.

In this study, we employed a TB mouse model using C3HeB/FeJ mice, which develop necrotizing granulomas upon *M. tuberculosis* infection and closely recapitulate the pathological features observed in human TB (7). Using scRNA-seq, we characterized the cellular composition and transcriptomic profiles of necrotizing granulomatous lesions in the lungs of this mouse model. The findings of this study will reveal unique transcriptional signatures of foamy macrophages, highlighting their potential as targets for novel TB diagnostics and host-directed therapeutics.

## Results

### Single cell transcriptomics reveals the cellular landscape of necrotizing granulomatous lesions

C3HeB/FeJ mice infected with *M. tuberculosis* via aerosol exposure developed necrotizing granulomatous lesions in the lungs at 12 weeks postinfection (p.i.) (Supplementary Figure 1), which is consistent with previous reports (30–34). We collected lung lesions with necrotic granulomas and prepared single-cell suspensions. To optimize the resolution of cellular transcriptomics, cell suspensions were further separated using Ficoll-Paque density gradient centrifugation, as necrotizing granulomas are primarily composed of abundant neutrophils and dead cell debris (30, 33, 35, 36). Cells collected from the interface layers of Ficoll-Paque solution were subjected to scRNA-seq (Figure 1A, B). Among 30,159 cells isolated from necrotizing granulomatous lesions, we manually annotated 11 major cell types. These included phagocytotic cells such as neutrophils and macrophages; dendritic cells (DCs) including conventional DCs (cDCs) and plasmacytoid DCs (pDCs); T cells including αβ T cell and γδ T cell subsets; natural killer (NK) cells; and B cells.

**Figure 1.**
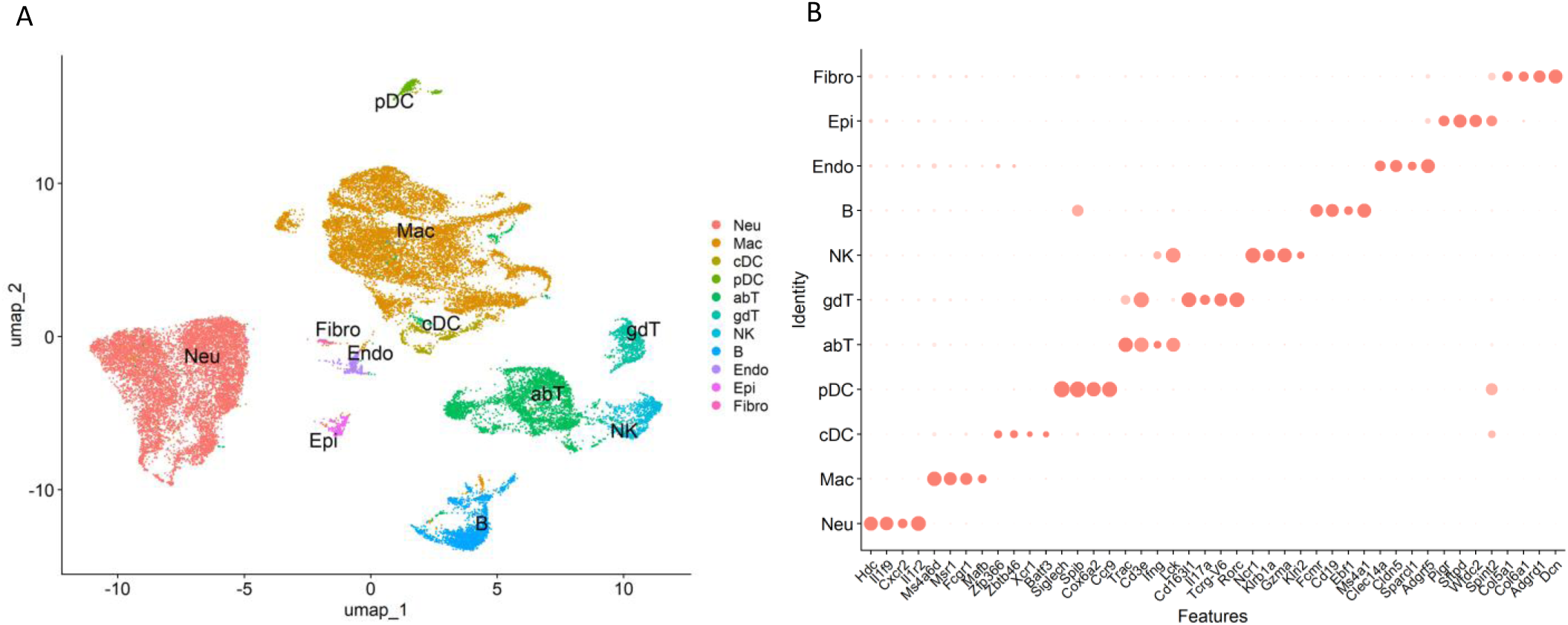
Single-cell transcriptomic landscape of necrotizing granulomatous lesions developed in *Mycobacterium tuberculosis*-infected lungs of C3HeB/FeJ mice. (A) Uniform manifold approximation and projection (UMAP) plot of 30,159 cells isolated from necrotizing granulomas, showing 11 major cell types. Neu, neutrophil; Mac, macrophage, cDC, conventional dendritic cell; pDC, plasmacytoid DC; abT, αβ T cell; gdT, γδ T cell; NK, natural killer cell; B, B cell; Endo, endothelial cell, Epi, epithelial cell, Fibro, fibroblast. (B) Dot plot showing the distinct expression of selected marker genes in each cell cluster.

Next, we assessed the subclusters of T cells within the necrotizing granulomatous lesions (Figure 2 and Supplementary Figure 2). Among CD4^+^ T cells, three clusters were identified: effector T cells, naïve T cells, and regulatory T cells (Tregs). For CD8^+^ T cells, two clusters were observed: cytotoxic and exhausted CD8^+^ T cells. Furthermore, we identified tissue-resident memory T cells (TRM) either expressing CD4 or CD8 within necrotizing granulomas.

**Figure 2.**
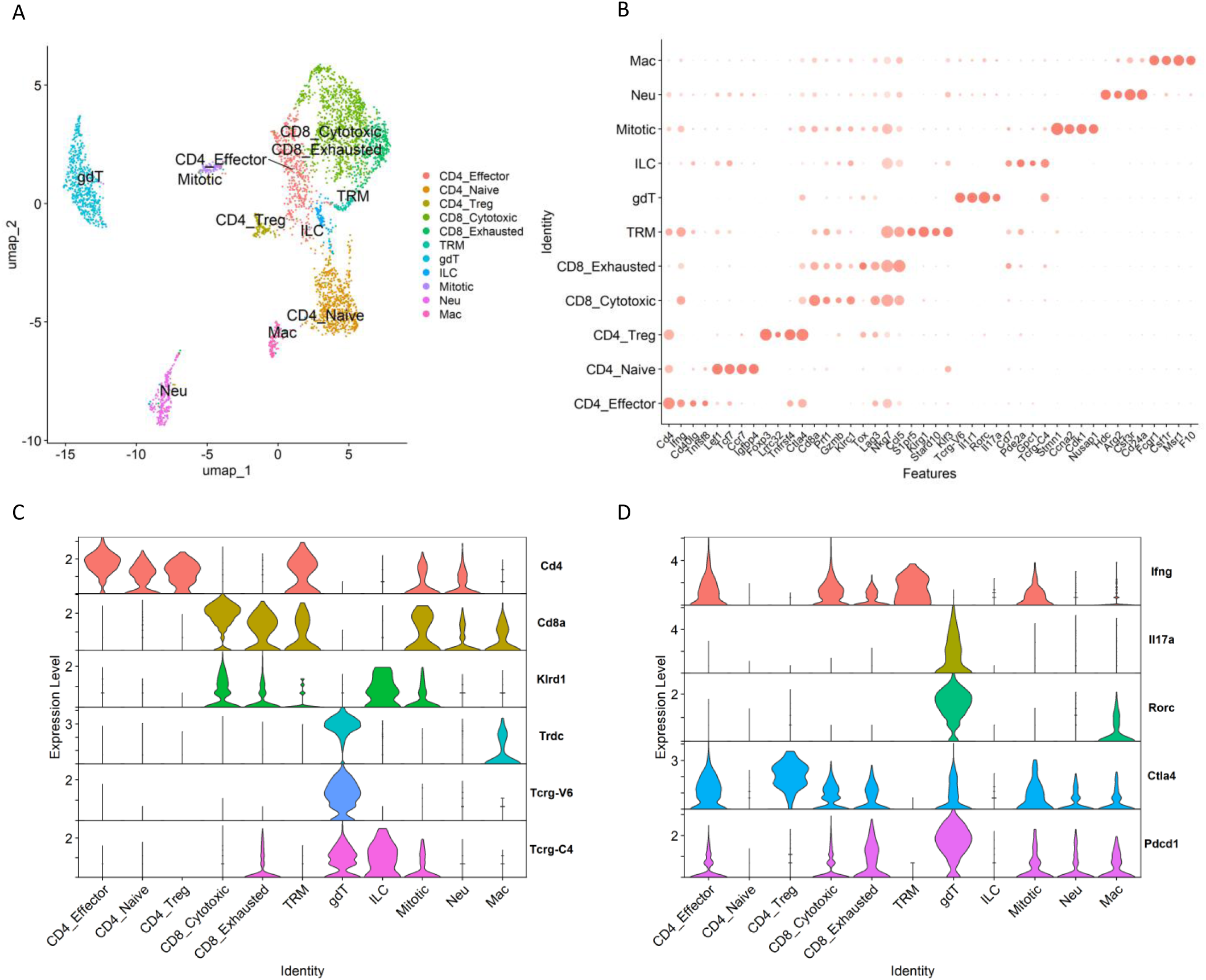
T cell clusters within necrotizing granulomatous lesions. T cell populations from necrotizing granulomas were further filtered to remove doubles and subsequently re-clustered. (A) UMAP plot showing T cell populations, including αβ T cells and γδ T cells. CD4_Effector, effector CD4^+^ T cell; CD4_Naive, naïve CD4^+^ T cell; CD4_Treg, regulatory CD4^+^ T cell; CD8_Cytotoxic, cytotoxic CD8^+^ T cell, CD8_exhausted, exhausted CD8^+^ T cell; TRM, tissue-resident memory T cell; gdT, γδ T cell; ILC, innate lymphoid cell; Mitotic, mitotic cell; Neu, neutrophil; Mac, macrophage. (B) Dot plot displaying the expression of selected marker genes for identified T cell clusters. (C, D) Violin plots displaying gene expression levels of T cell receptors and co-receptor molecules (C) and cytokines and immune checkpoint molecules (D).

During *M. tuberculosis* infection, γδ T cells participate in the immune response in the lungs (37). In particular, IL-17-mediated immunity is critical for γδ T cells to perform their function in host defense in TB murine models (38). In necrotizing granulomas, γδ T cells expressing *Il17a* were identified (Figure 2C, D). These γδ T cells also expressed *Pdcd1*, which encodes programmed cell death-1 (PD-1), an immune checkpoint molecule inhibiting immune responses (39). These results imply that *Pdcd1^+^* γδ T cells contribute to the pathogenesis of necrotizing granulomas in the lungs of *M. tuberculosis*-infected C3HeB/FeJ mice. Among other T cell types, we identified a unique population co-expressing *Tcrg* (T cell receptor gamma) and *Klrd1* (encoding an NK cell receptor) (Figure 2). This population also expressed *Gata3*, suggesting that they resemble type 2 innate lymphoid cells (ILC2). Following the removal of doublet cells from scRNA-seq data, clusters expressing characteristic genes of neutrophils or macrophages remained, suggesting physiological interactions between T cells and these phagocytotic cells.

### Macrophage populations in necrotizing granulomatous lesions

To investigate the hallmark of macrophage populations within necrotizing granulomas, we analyzed their gene expression profiles. We subdivided the macrophage populations into 29 clusters and explored the characteristic of each cluster (Supplementary Figure 3A). Given that *Plin2* expression is significantly upregulated in foamy macrophages within necrotizing granulomas (29, 30), we identified the clusters exhibiting significantly elevated *Plin2* expression compared to other clusters (Supplementary Figure 3B). As expected, *Plin2-*expressing clusters were localized in close proximity to one another (Figure 3A). To characterize the remaining clusters, we analyzed their gene expression profiles and found that macrophages expressing *Cxcl10*, *Clec4e*, *C1qc*, *Ighm*, *S100a9*, or *Siglecf* also represented distinct macrophage populations within the necrotizing granulomatous lesions (Figure 3A and Supplementary Figure 3C). Moreover, we confirmed that the expression levels of the corresponding genes in their respective clusters were significantly higher than those in other clusters (Figure 3B).

**Figure 3.**
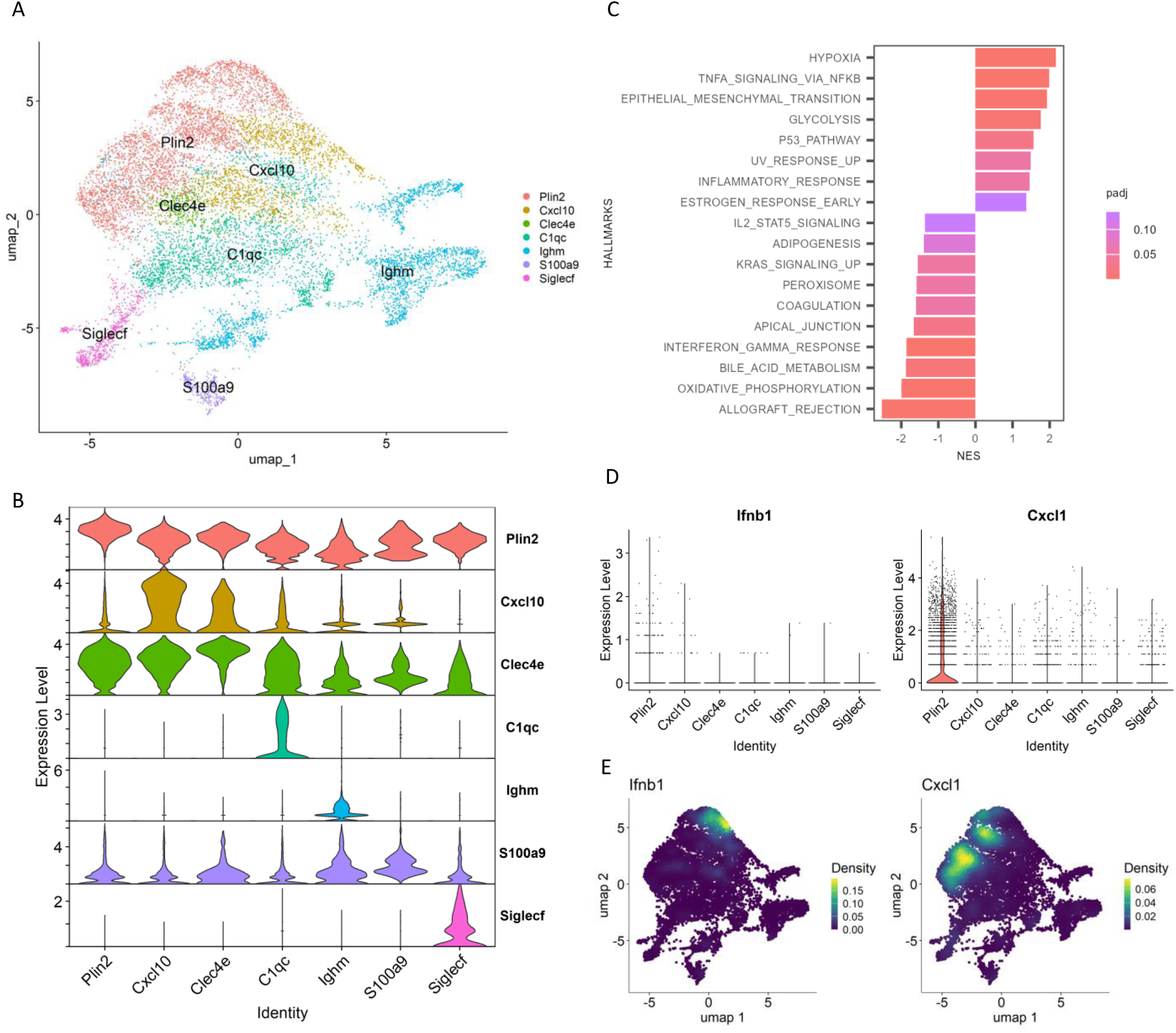
Identification of foamy macrophage associated subclusters. Macrophage populations in necrotizing granulomas were re-clustered and analyzed. (A) UMAP plot of macrophage clusters in necrotizing granulomatous lesions, characterized by signature genes. (B) Violin plot showing significantly elevated expression of the indicated genes in specific clusters. (C) Gene set enrichment analysis (GSEA) of *Plin2*-expressing macrophage cluster. Significantly enriched hallmarks are depicted, representing both activated and suppressed hallmarks in the *Plin2*+ cluster compared with other clusters. (D, E) Expression profiles of genes involved in tuberculosis (TB) exacerbation in macrophages. Violin (D) and UMAP (E) plots demonstrating g *Ifnb1* and *Cxcl1* expression in macrophage clusters.

To further investigate the gene expression profile of the *Plin2*-expressing (*Plin2*^+^) cluster, we performed gene set enrichment analysis (GSEA)

(Figure 3C). GSEA revealed that genes associated with hypoxia, TNF-α signaling, glycolysis, and inflammatory responses were upregulated, whereas those associated with oxidative phosphorylation and type II interferon (IFN) responses were downregulated in the *Plin2*^+^ cluster compared to other macrophage clusters. Further, we also performed Weighted Gene Coexpression Network Analysis (WGCNA) to construct a gene network within the macrophage populations from necrotizing granulomas (Supplementary Figure 4). Correlation analysis of module–trait relationship revealed the presence of one module exhibiting strongly positive correlation with the *Plin2*^+^ cluster. Genes within this module were specifically and highly expressed in the *Plin2*^+^ cluster. Gene Ontology (GO) enrichment analysis demonstrated that these genes are involved in glycolysis, hypoxia, TNF signaling pathway, and lipid metabolism and atherosclerosis.

We investigated the gene expression profiles of additional macrophage clusters within necrotizing granulomatous lesions using GSEA (Supplementary Figure 5). Clusters expressing *Cxcl10 or Clec4e* exhibited gene expression signatures characteristic of pro-inflammatory macrophages. In contrast, clusters expressing *C1qc*, *Ighm*, or *Siglecf* showed gene expression profiles associated with anti-inflammatory macrophages. A more detailed analysis revealed that *Ighm*^+^ macrophages expressed several anti-inflammatory genes, including *Cd244a*, *Nr4a1*, *Clec4a1*, and *Clec4a2* (Supplementary Figure 6A). Since *Siglecf* is a well-established marker of alveolar macrophages (40), the *Siglecf*^+^ cluster was identified as alveolar macrophages. *Siglecf-*expressing macrophages expressed *Alox5*, a gene that promotes anti-inflammatory polarization and contributes to increased susceptibility to *M. tuberculosis* infection (41, 42) (Supplementary Figure 6B).

### Novel polarization of *Plin2*-expressing macrophages

*Nos2* and *Arg1*, well-established markers of macrophage polarization, play key roles in regulating immune responses (43, 44). We have previously demonstrated that Plin2-expressing macrophages express Nos2 or Arg1 in necrotizing granulomas (30), suggesting that the polarization of foamy macrophages exhibits either pro-inflammatory or anti-inflammatory characteristics. Accordingly, scRNA-seq profiling revealed that *Nos2* was highly expressed in pro-inflammatory macrophage clusters including the *Plin2*^+^ cluster (Figure 4A). Moreover, *Arg1* expression was predominantly observed in the *Plin2*^+^ cluster. Feature plots revealed the presence of *Plin2*^+^ macrophages co-expressing *Nos2* and *Arg1* (Figure 4B). Further, we investigated the transcriptional factors associated with the regulation of *Nos2* or *Arg1* expression in macrophages within necrotizing granulomas. Among them, we found that *Irf7* expression was correlated with *Nos2* expression patterns in macrophage clusters (Figure 4A). *Irf7* regulates macrophage polarization toward both pro-inflammatory and anti-inflammatory states (45). *Fosl1* encodes a transcriptional factor promoting pro-inflammatory polarization by repressing *Arg1* expression (46). We found *Fosl1* expression in a subset of *Arg1*^+^ macrophages within the *Plin2*^+^ cluster (Figure 4C). These results suggest that *Irf7* and *Fosl1* are associated with the polarization dynamics of *Plin2^+^* macrophages in necrotizing granulomas.

**Figure 4.**
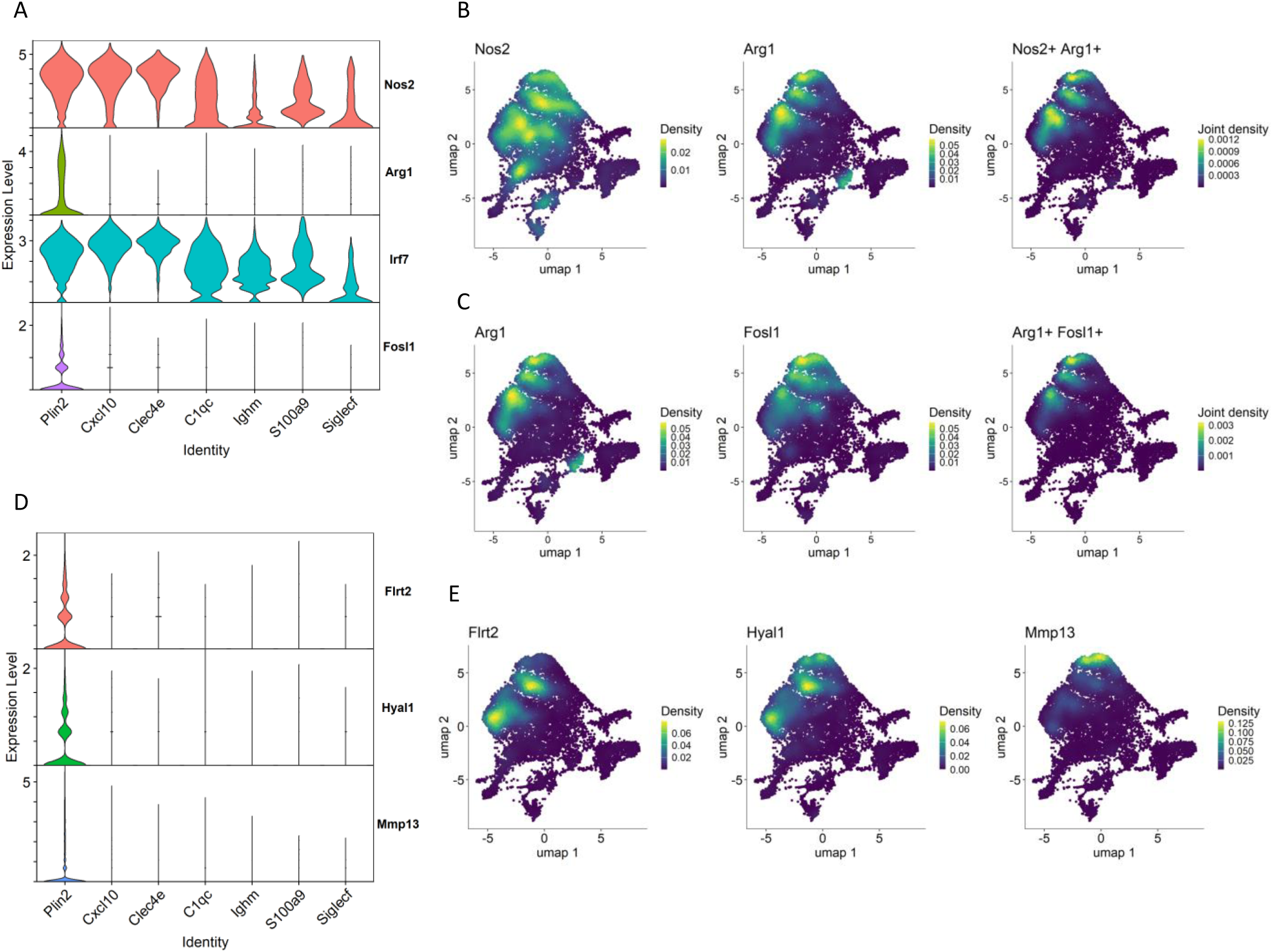
Novel polarization and gene expression profiles of the *Plin2^+^* macrophage cluster. (A-C) Novel polarization of the *Plin2+* cluster. (A) Violin plots showing the gene expression of *Nos2* and *Arg1* and their transcriptional factors *Irf7* and *Fosl1* in macrophage clusters. (B) UMAP plot illustrating expression and co-expression of *Nos2* and *Arg1* in macrophages. (C) Expression and co-expression of *Arg1* and *Fosl1* in macrophages. (D, E) Novel gene expression profiles in the *Plin2+* cluster. (D) Violin plots demonstrating the expression of *Flrt2*, *Hyal1*, and *Mmp13*, specifically expressed in the *Plin2*+ cluster. (E) UMAP plots demonstrating the expression of *Flrt2*, *Hyal1*, and *Mmp13*, specifically associated with the *Plin2^+^* cluster.

We further analyzed polarization states of macrophages derived from myeloid cell lineages in the whole lungs of *M. tuberculosis*-infected *Sp140* knockout (KO) mice, using the data from Kotov et al. (28). In C3HeB/FeJ mice, the reduced gene expression of both *Sp110* and *Sp140* in *Sst1* locus contributes to increased susceptibility to *M. tuberculosis* infection (47–49). In addition, similar to C3HeB/FeJ mice, *Sp140* KO mice also exhibit increased susceptibility to *M. tuberculosis* infection, indicating that *Sp140* is a key determinant of host vulnerability to the infection (50). Macrophage populations derived from *M. tuberculosis*-infected *Sp140* KO mice were re-clustered based on the original annotations (Figure 5A). Subsequently, *Plin2* expression was found to be elevated in the cluster of interferon-stimulated gene-positive (ISG^+^) interstitial macrophages (IMs) compared to other macrophage clusters. This result suggests that ISG^+^ IMs correspond to the *Plin2*^+^ cluster identified in our study (Figure 5B). This was further supported by an integrated analysis of macrophage populations from C3HeB/FeJ mice and *Sp140* KO mice (Supplementary Figure 7A and B). Notably, *Nos2* was expressed in ISG^+^ IM and IM populations, whereas *Arg1* was dominantly expressed in ISGs^+^ IMs (Figure 5B and Supplementary Figure 7C). Moreover, macrophages co-expressing *Nos2* and *Arg1* were observed in a subset of the ISG^+^ IM cluster (Figure 5C).

**Figure 5.**
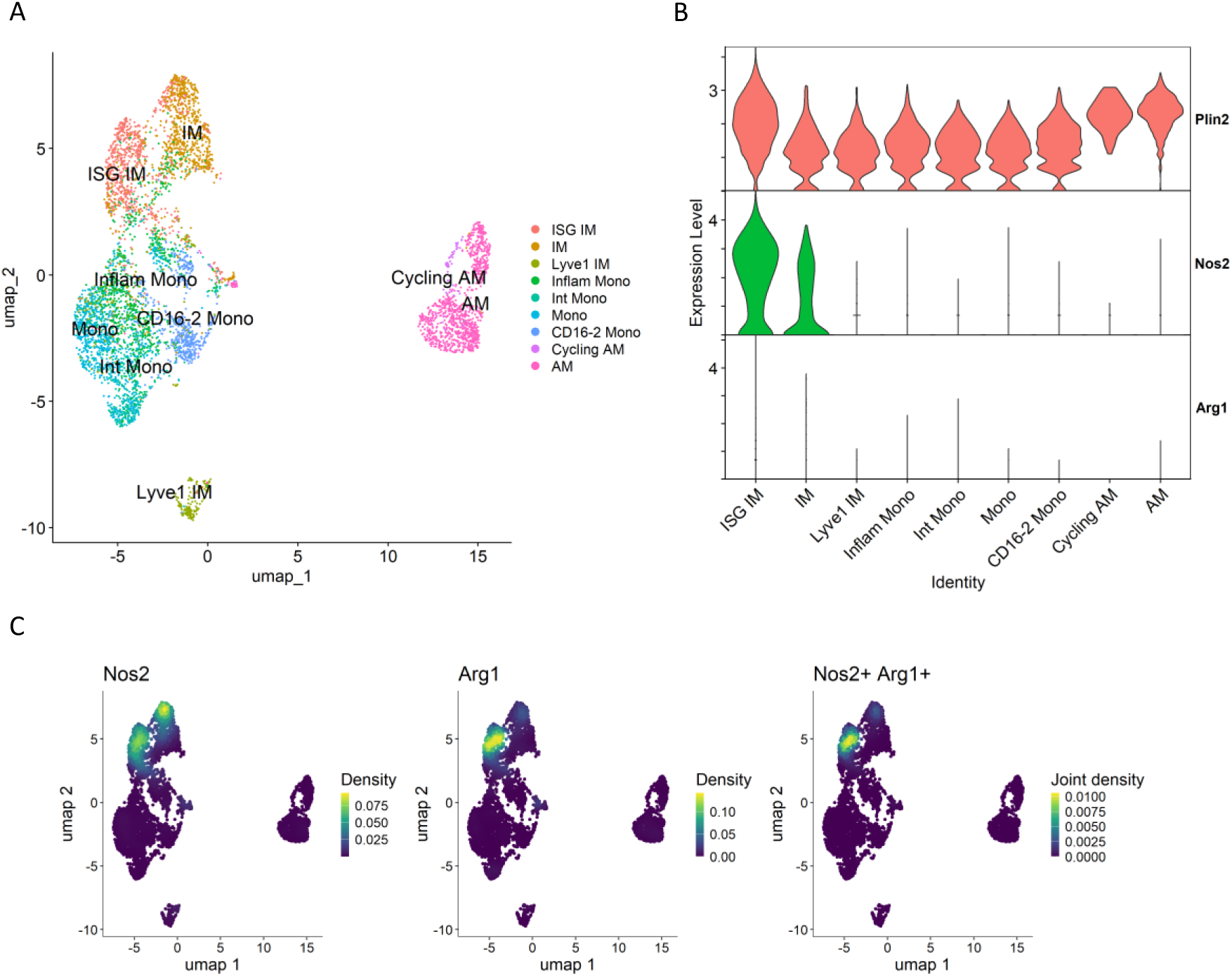
Macrophages derived from *M. tuberculosis*-infected *Sp140*-deficient mouse lungs. Macrophage populations of the GSE216023 data from Kotov et al. (28) were analyzed. (A) UMAP plot of macrophage populations from *Sp140* knockout (KO) mouse lungs infected with *M. tuberculosis*. (B) Volin plot showing expression levels of *Plin2*, *Nos2* and *Arg1* in macrophage clusters derived from *Sp140* KO mice. (C) UMAP plot showing expression and co-expression of *Nos2* and *Arg1* in macrophages derived from *Sp140* KO mice.

These results suggest the emergency of a novel macrophage polarization state in ISG ^+^ IMs derived from *M. tuberculosis*-infected *Sp140* KO mice, corresponding to the *Plin2*^+^ cluster of C3HeB/FeJ mice identified in our study.

### Gene expression characteristic of *Plin2*-expressing macrophages

Type I IFNs and neutrophils contribute to TB exacerbation (35, 51). Accordingly, we investigated the expression of *Ifnb1*, a type I IFN gene, and *Cxcl1*, a chemokine gene involved in neutrophil recruitment, in macrophage populations (Figure 3D, E). Both genes were specifically expressed in the *Plin2*^+^ cluster, consistent with previous reports (28, 35). pDCs regulate viral infections by producing a large amount of type I IFNs (52). In the context of *M. tuberculosis* infection, the depletion of pDCs results in increased bacterial burdens in the lungs (28). Lee et al. demonstrated that *M. tuberculosis*-infected neutrophils can stimulate pDCs to produce type I IFNs (53). Therefore, we assessed the expression levels of *Ifnb1* in pDCs (Figure 6). However, *Ifnb1* expression were predominantly detected in macrophages rather than pDCs. In contrast, pDCs expressed *Il34* and *Kmo*. *Il34* encodes an anti-inflammatory cytokine that regulates the expression of pro-inflammatory cytokines and promotes macrophage polarization toward an anti-inflammatory phenotype (54). *Kmo* encodes kynurenine 3-monooxygenase, which catalyzes the conversion of kynurenine to 3-hydroxykynurenine (3-HK), a metabolite inducing cellular damage and apoptosis via oxidative stress (55). These results suggest that pDCs are localized to necrotizing granulomatous lesions and contribute to the pathogenicity of *M. tuberculosis* infection via the expression of immunomodulatory and cytotoxic factors.

**Figure 6.**
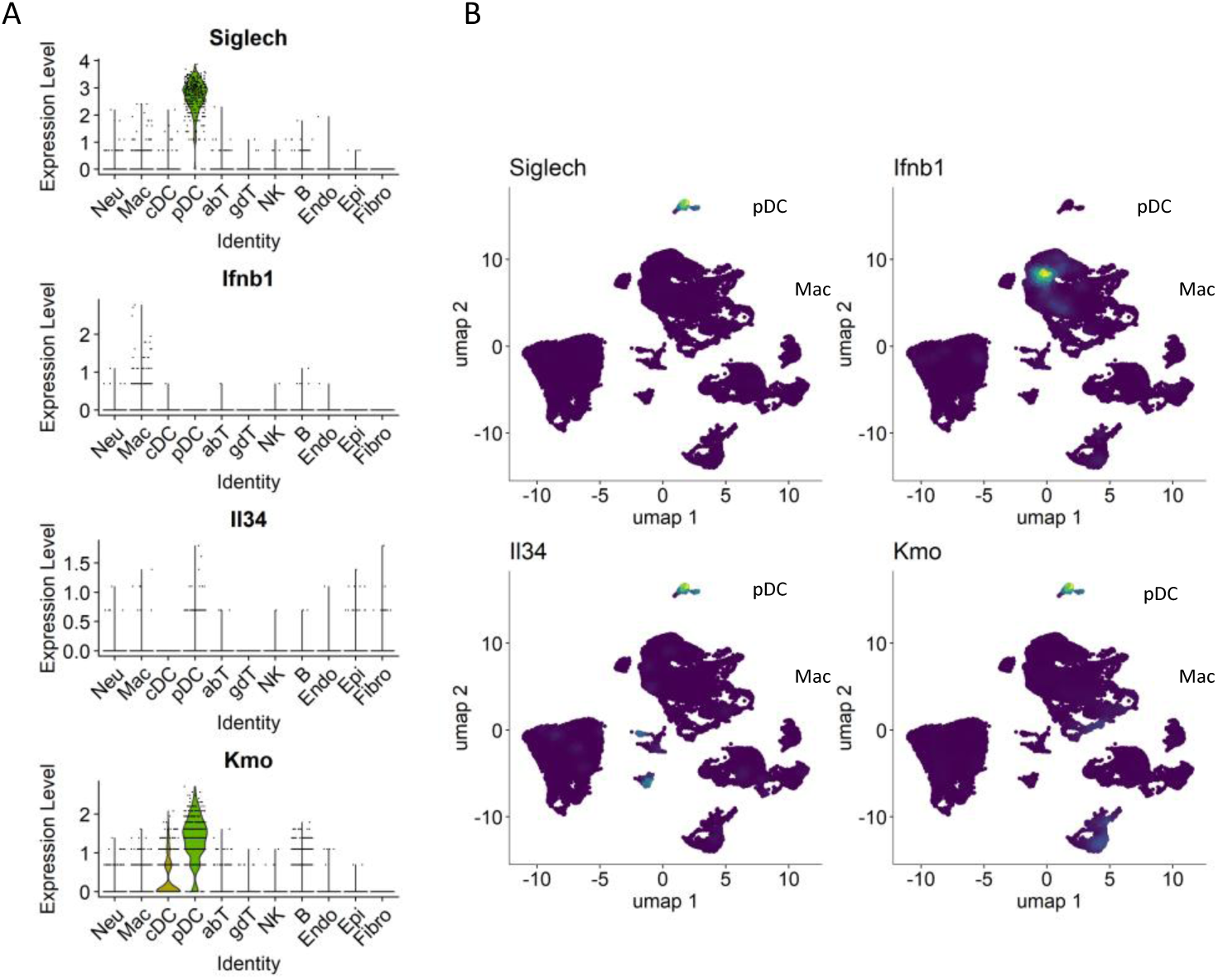
Expression of *Il34* and *Kmo* in necrotizing granulomas of *M. tuberculosis*-infected C3HeB/FeJ mice. Violin (A) and UMAP (B) plots showing the gene expression of *Ifnb1*, *Il34*, *Kmo* and *Siglech* (pDC marker) in necrotizing granulomas derived from *M. tuberculosis*-infected C3HeB/FeJ mice.

We investigated novel gene expression signatures characteristic of the *Plin2*^+^ cluster within necrotizing granulomas. Among the genes expressed in the *Plin2*^+^ cluster, *Flrt2*, *Hyal1*, and *Mmp13* were selected for further examination, due to their specific and enriched expression patterns (Figure 4D, E). IHC revealed that these proteins were localized to the rim regions of necrotizing granulomas, consistent with Plin2 localization (Figure 7A). Immunofluorescence microscopy (IFM) showed co-localization of all the three proteins with Plin2^+^ cells. Notably, the fluorescent signals of Flrt2, Hyal1, and Mmp13 displayed distinct subcellular localization patterns, whereas Plin2 was predominantly localized to the cytosolic regions of the same vacuolated cells (Figure 7B). These results suggest that *Flrt2*, *Hyal1*, and *Mmp13* are novel molecular markers characterizing *Plin2*^+^ macrophages in necrotizing granulomas.

**Figure 7.**
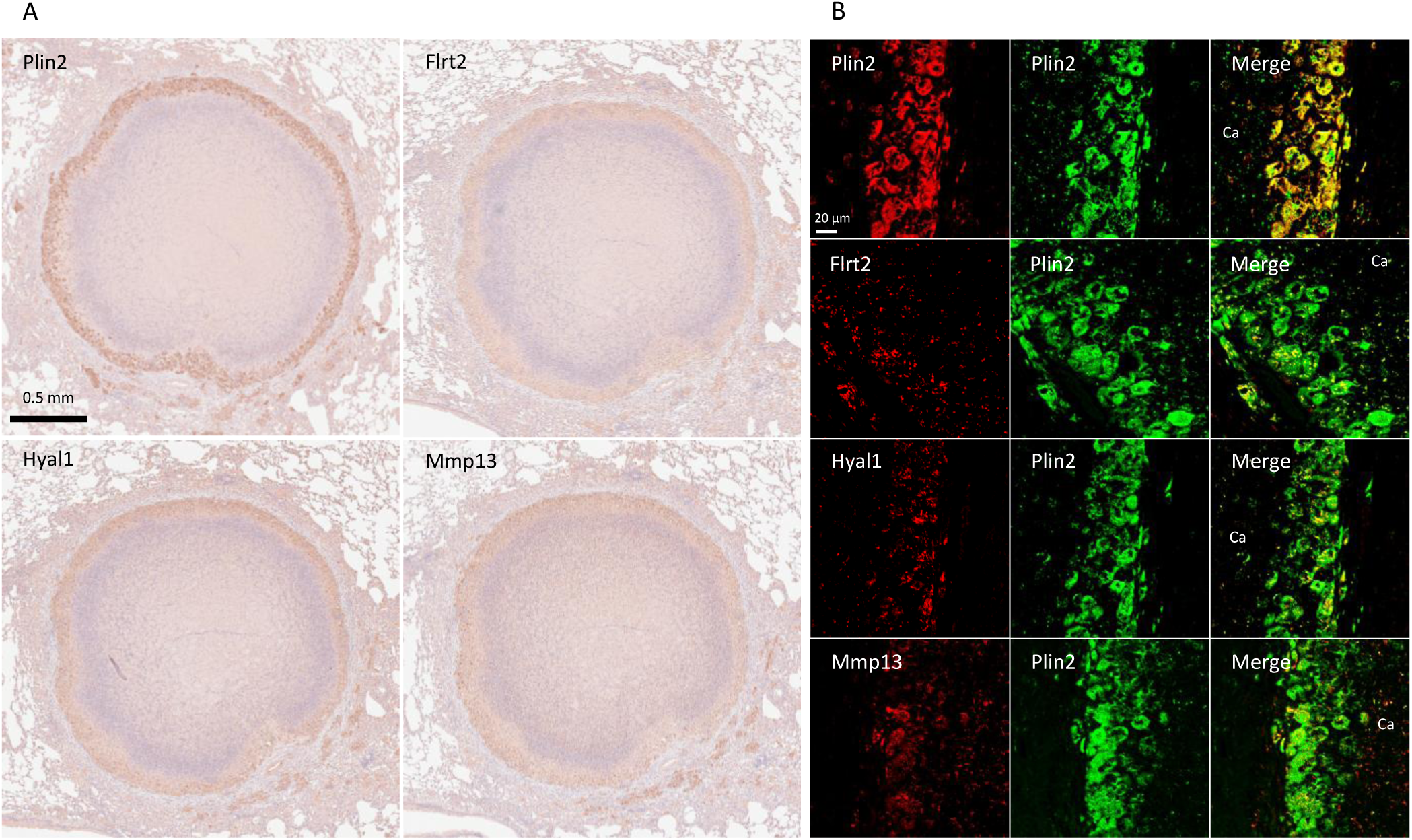
Localization of the novel signature of foamy macrophages in necrotizing granulomas. (A) Immunohistochemistry images showing the localization of Flrt2, Hyal1, and Mmp13 in necrotizing granulomas. The marker for foamy macrophages, Plin2 is also shown. (B) Immunofluorescence microscopic images demonstrating co-localization of Plin2 with the indicated proteins in necrotizing granulomas. Ca, caseous necrosis region within necrotizing granulomas.

## Discussion

scRNA-seq has been widely utilized to evaluate the cellular transcriptomics of tissues or blood samples from TB patients, as well as from *M. tuberculosis*-infected non-human primates, mice and other experimental models (56) However, the detailed cellular composition of necrotizing granulomatous lesions, a hallmark of TB pathology (5, 7), remains incompletely characterized.

In this study, we isolated single-cell suspensions from necrotizing granulomatous lesions developed in the lungs of *M. tuberculosis*-infected C3HeB/FeJ mice, and performed scRNA-seq to comprehensively investigate their cellular landscape. We identified 11 major cell type, including immune cells such as neutrophils, macrophages, dendritic cells, T cells, NK cell, and B cells (Figure 1). To prepare single-cell suspensions from necrotizing granulomatous lesions, we performed Ficoll-Paque density gradient centrifugation to reduce the number of dead cells and neutrophils. However, due to the heterogeneity and activation states of neutrophils, their density can vary, resulting in partial retention at the interface of the Ficoll-Paque gradient.

Among T cells, CD4^+^ T cells, particularly central memory *Mtb*-specific CD4^+^ T cells, play a crucial role in protective immunity against *Mtb* infection (56). In this study, we identified CD4^+^ T cells in necrotizing granulomatous lesions as naïve, effector or TRM types (Figure 2 and Supplementary Figure 2B). However, central memory CD4^+^ T cells (*Cd44*^+^, *Sell*^+^, and *Ccr7*^+^) were not identified, suggesting that these cells do not localize to necrotizing granulomatous lesions. Naïve CD4^+^ T cells were enriched in the lesions, consistent with previous reports in non-human primate and murine models of *M. tuberculosis* infection (25–27). Moreover, *Pdcd1*^+^ γδ T cells were identified within necrotizing granulomatous lesions (Figure 2). Although IL-17-producing γδ T cells play a protective role in the lungs of *M. tuberculosis*-infected mice (58), the presence of *Pdcd1*^+^ γδ T cells has not been previously reported in this context. In the experimental autoimmune encephalomyelitis mouse model, *Pdcd1*^+^ γδ T cells have been implicated in promoting disease pathogenesis (59). These findings suggest that *Pdcd1*^+^ γδ T cells may similarly contribute to the immunopathology of necrotizing granulomas during *M. tuberculosis* infection.

In macrophage populations within necrotizing granulomas, the *Plin2^+^* cluster was identified based on *Plin2* expression in the clusters (Figure 3). Because Plin2 is the marker of foamy macrophages in the lungs of TB patients and *M. tuberculosis*-infected C3HeB/FeJ mice (29, 30), we referred to the *Plin2^+^* cluster as the foamy macrophage population. Furthermore, GSEA and WGCNA revealed that gene expression profiles of the *Plin2*^+^ cluster were consistent with the characteristics of foamy macrophages in necrotizing granulomas (29, 30).

Previously, we have demonstrated that foamy macrophages in necrotizing granulomas express typical macrophage polarization markers, either *Nos2* or *Arg1*, suggesting their differentiation into a pro-inflammatory or anti-inflammatory state (30). scRNA-seq revealed that *Plin2^+^* macrophages express *Nos2* or *Arg1* (Figure 4). Moreover, we identified a subset of *Plin2^+^* macrophages co-expressing both *Nos2* and *Arg1*, suggesting a novel polarization state of foamy macrophages in necrotizing granulomas. A similar polarization profile was also observed in macrophage populations isolated from the lungs of *M. tuberculosis*-infected *Sp140* KO mice (28) (Figure 5). Foamy macrophages exhibiting the dual expression profile may represent a transitional state between pro-and anti-inflammatory phenotypes. To investigate this possibility, we performed an integrated analysis of macrophage populations from both C3HeB/FeJ and *Sp140* KO mice (Supplementary Figure 7). This analysis revealed a distinct correspondence between the *Plin2^+^* cluster in C3HeB/FeJ mice and ISG^+^ IMs in *Sp140* KO mice, as well as between the *C1qc+* cluster and IMs, the *Ighm+* cluster and monocyte clusters, the *Siglecf^+^* cluster and alveolar macrophage clusters. Macrophage differentiation within necrotizing granulomas was investigated using trajectory analysis (Supplementary Figure 8). We defined cluster number 11, corresponding to monocytes and intermediate monocytes in Supplementary Figures 3A and 6A as the start point for differentiation. The trajectory analysis suggested that the *Igmh^+^* cluster differentiates into *Cxcl10*-expressing or *Clec4e*-expressing macrophages. Subsets of these macrophages subsequently transitioned into foamy macrophages. Together, these results suggest that monocytes infiltrating infected lungs differentiate into IMs and ISG^+^ IMs, subsets of which further progress toward a foamy macrophage phenotype.

Type I IFNs and neutrophils are key exacerbating factors in TB, contributing to disease progression by promoting inflammation and inducing the release of neutrophil extracellular traps (NETs) (35, 50, 51). In particular, neutrophils and their remnants resulting from NETosis accumulate in the caseous necrosis regions of necrotizing granulomas. We found a subset of *Plin2^+^* macrophages expressing *Ifnb1* or *Cxcl1* (Figure 3D, E). scRNA-seq revealed that ISG^+^ IMs and IMs express type I IFNs in the lungs of *M. tuberculosis*-infected *Sp140* KO mice (28). Chowdhury et al. demonstrated that type I IFNs are primarily expressed by epithelioid cells proximal to the caseous necrosis regions in necrotizing granulomas developed in C3HeB/FeJ mice and non-human primate models (35), which is consistent with our findings.

Moreover, Chowdhury et al. demonstrated that pDCs within necrotizing granulomatous lesions are not significantly associated with type I IFN expression. In agreement with this, our study revealed that pDCs within necrotizing granulomatous lesions do not predominantly express *Ifnb1*, but rather express other immunosuppressive genes, such as *Il34* and *Kmo* (Figure 6). These results suggest that Cxcl1, secreted by foamy macrophages in necrotizing granulomas, contribute to the recruitment of neutrophils into the caseous necrosis regions, where NETosis may be subsequently induced by type I IFNs.

We investigated unique molecular signatures of the *Plin2+* cluster in necrotizing granulomas. We found that three proteins, Flrt2, Hyal1, and Mmp13, were specifically localized to Plin2^+^ macrophages (Figures 4 and 7). *Flrt2* regulates macrophage differentiation and activate Akt/mTOR signaling (60). Given the involvement of mTORC1 signaling in the differentiation of foamy macrophages during *M. tuberculosis* infection (61), *Flrt2* may contribute to foamy macrophage differentiation in necrotizing granulomas. *Hyal1* encodes a lysosomal hyaluronidase that digests extracellular matrix (62). We found that a subset of *Cxcl1*-expressing *Plin2^+^* macrophages also expressed *Hyal1* (Supplementary Figure 6C), suggesting that *Hyal1* facilitates neutrophil recruitment into the caseous necrosis regions. *Mmp13* encodes a metalloprotease involved in collagen remodeling in atherosclerotic plaques (63). Foamy macrophages exhibit a characteristic arrangement around the necrotic core within necrotizing granulomas, suggesting that Mmp13 expressed by foamy macrophages may regulate cell–cell interaction or tissue remodeling.

Taken together, these identified molecules are involved in the development of necrotizing granulomas through the regulation of foamy macrophage differentiation, localization, and function.

In conclusion, we conducted an in-depth single-cell transcriptomic analysis of necrotizing granulomatous lesions using a TB mouse model that closely recapitulates the pathological features observed in TB patients. Our results revealed novel cellular signatures within necrotizing granulomas, with particular emphasis on the characterization of foamy macrophage. These insights into the cellular and molecular landscape of necrotizing granulomas advance our understanding of TB pathogenesis and will facilitate the development of novel diagnostic tool and host-directed therapeutic drugs for TB.

## Materials and Methods

### Ethics statement

All animal experiments in this study were approved by the Animal Care and Use Committee of The Research Institute of Tuberculosis (RIT) (permission number ID 2022-02) and conducted in accordance with the RIT ethical guidelines for animal care and use.

### Mouse model and infection

C3HeB/FeJ mice were purchased from Jackson Laboratory and housed in a filtered-air, laminar-flow cabinet under specific pathogen-free conditions at the animal facility of the RIT. Mice were provides with sterile bedding, water, and mouse chow. Specific pathogen-free status was verified by monitoring sentinel mice housed within the colony. Mice aged 6–10 weeks were transferred to the biosafety level III animal facility of the RIT. For *M. tuberculosis* infection, frozen stocks of the *M. tuberculosi*s Erdman strain stored at-80°C was used as previously described (64). Mice were infected with approximately 100 CFU of *M. tuberculosis* bacilli via aerosol using an infection exposure system (Glas-Col).

### Preparation of single cells from necrotizing granulomatous lesions

At 12 weeks p.i., infected mice were euthanized by exsanguination under anesthesia with 0.75 mg/kg medetomidine, 4.0 mg/kg midazolam, and 5.0 mg/kg butorphanol via the intraperitoneal route. The lungs were excised and subsequently dissected to collect infected lesions including necrotizing granulomas (Supplementary Figure 1). For the preparation of single-cell suspensions, infected lungs from three or four mice were pooled to collect more than 10 lesions containing necrotizing granulomas. Necrotizing granulomatous lesions were minced and incubated in a collagenase/hyaluronidase/DNase I solution (Stemcell) with RNase inhibitor at 0.2 U/μL. Following red blood cell lysis, the dissociated cells were resuspended in PBS containing 2% fetal bovine serum and RNase inhibitor at 0.2 U/μL. To remove dead cells and a large proportion of neutrophils, cell suspensions were subjected to Ficoll-Paque density gradient centrifugation, followed by the collection of cells from the interface layer according to the manufacturer’s instructions (Cytiva).

### Construction of scRNA-seq libraries and sequencing

Isolated cell suspensions from necrotizing granulomatous lesions were subjected to scRNA-seq library constructions using Chromium Fixed RNA Profiling Reagent Kit (10x Genomics). Cells were fixed with a fixation solution containing formaldehyde for 24 h at 4°C. Inactivation of *M. tuberculosis* in the fixed samples was confirmed by CFU assay. Fixed cells were further processed to construct scRNA-seq libraries according to the manufacturer’s protocol. The resulting libraries were sequenced on NextSeq 1000 (Illumina).

## Data analysis

Raw sequencing reads were aligned against the mouse reference genome (mm10) using Cellranger version 7.0.1 (10x Genomics). Subsequent analyses were performed in R version 4.4.1 using the Seurat package version 5.2.1 (65). Data from four independent samples were filtered to include cells with 500–6000 genes and less than 20% mitochondrial reads. Data were normalized using the NormalizeData function, followed by the identification of highly variable features using the FindVariableFeatures function with the variance stabilizing transformation method, selecting the top 2000 most variable genes. Principal component analysis (PCA) was performed using the RunPCA function. To correct for batch effects, the Harmony algorithm was applied using the RunHarmony function. Following batch correction, uniform manifold approximation and projection (UMAP) was computed using the first 30 dimensions. A shared nearest neighbor (SNN) graph was then constructed using the FindNeighbors function with the same set of dimensions. Clustering was performed using the FindClusters function with a resolution parameter set to 0.7. The resulting dataset was processed with scDblFinder version 1.18.0 (66) to remove doublet cells. The filtrated data were reprocessed through data scaling, PCA, UMAP, followed by the construction of SNN graph and clustering.

Cell clusters were manually annotated based on specific gene expression patterns of the respective cell types.

Gene expression analysis, GSEA, WGCNA, GO analysis, and trajectory analysis were performed using Nebulosa version 1.14.0 (67), fGSEA version 1.30.00 (68), WGCNA version 1.73 (69), ShinyGO version 0.82 (70), and Monocle3 version 1.3.7 (71), respectively.

### Immunohistochemistry and immunofluorescence microscopy

IHC and IFM were performed as previously described (30, 36).

Antibodies used in this study are listed in Supplementary Table S1. IHC and IFM samples were visualized using a NanoZoomer S60 (Hamamatsu Photonics) and a FV4000 (Evident), respectively.

## ACKNOWLEDGMENTS

This study was supported by the Emerging/Re-emerging Infectious Diseases Project of the Japan Agency for Medical Research and Development (25fk0108673, 25fk0108674, 25gm1610013, 25wm0225028, 25fk0108703, 25fk0108730), and Grants-in-Aid for Scientific Research, Japan Society for the Promotion of Science (20KK0197, 22K07065).

We thank Ms. Miyako Seto and Ms. Mariko Ogasawara in Department of Pathophysiology and Host Defense for technical supports.

SS, MH, NK designed the project and analyzed data, and wrote and revised the manuscript. SS, SO, HN, performed experiments. All authors approved the manuscript.

The authors declare that the research was conducted in the absence of any commercial or financial relationships that could be construed as a potential conflict of interest.

